# The material properties of naked mole-rat hyaluronan

**DOI:** 10.1101/447631

**Authors:** Yavuz Kulaberoglu, Bharat Bhushan, Fazal Hadi, Sampurna Chakrabarti, Walid T. Khaled, Kenneth S. Rankin, Ewan^l^ St. John Smith, Daniel Frankel

**Affiliations:** Department of Pharmacology, University of Cambridge, Tennis Court Road, Cambridge, CB2 1PD, UK; Northern Institute for Cancer Research, Medical School, Newcastle University, Paul O’Gorman Building, Framlington Place, Newcastle upon Tyne, NE2 4HH, UK; School of Engineering, Newcastle University, Merz Court, Newcastle upon Tyne NE1 7RU, UK

## Abstract

Hyaluronan (HA) is a key component of the extracellular matrix. Given the fundamental role of HA in the cancer resistance of the naked mole-rat (NMR), we undertook to explore the structural and soft matter properties of this species-specific variant, a necessary step for its development as a biomaterial. We examined HA extracted from NMR brain, lung, and skin, as well as that isolated from the medium of immortalised cells. In common with mouse HA, NMR HA forms a range of assemblies corresponding to a wide distribution of molecular weights. However, unique to the NMR, are highly folded structures, whose characteristic morphology is dependent on the tissue type. Skin HA forms tightly packed assemblies that have spring-like mechanical properties in addition to a strong affinity for water. Brain HA forms three dimensional folded structures similar to the macroscopic appearance of the gyri and sulci of the human brain. Lung HA forms an impenetrable mesh of interwoven folds in a morphology that can only be described as resembling a snowman. Unlike HA that is commercially available, NMR HA readily forms robust gels without the need for chemical cross-linking contrasting. NMR HA gels sharply transition from viscoelastic to elastic like properties upon dehydration or repeated loading. In addition, NMR HA can form ordered thin films with an underlying semi-crystalline structure. Given the role of HA in maintaining hydration in the skin it is plausible that the folded structures contribute to both the elasticity and youthfulness of NMR skin. It is also possible that such densely folded materials could present a considerable barrier to cell invasion throughout the tissues, a useful characteristic for a biomaterial.

---

Hyaluronan (HA), also known^1,2^ as hyaluronic acid, is an extracellular matrix (ECM) polymer found in most tissues. Three key enzymes are involved in HA synthesis, HA synthase 1, 2 and 3 (HAS1-3), and it is subsequently broken down by hyaluronidase^3^. The regulation of HA turnover is highly important with there being a clear link between HA metabolism and cancer progression, an abundance of HA being an indicator of poor patient prognosis for a number of tumours^4–8^. HA’s role in disease progression is perhaps unsurprising given its function in cell-ECM interactions^9–11^. The main HA receptor is CD44, a cell surface adhesion receptor that binds a range of ligands and is itself associated with metastasis^12^. The biological function of HA is often related to its molecular weight with a range of polymer sizes found depending on tissue of origin^13,14^. Given the importance of ECM-cell interactions in disease progression and HA-cell interaction in particular, it would be intriguing to examine the structure/assembly of HA in a cancer resistant species.

The naked mole-rat (NMR, *Heterocephalus glaber*) has a remarkable biology. They can live for up to ten times longer than similarly sized rodents and rarely develop cancer^15–17^. In 2013, a paper reported that HA contributed to the NMR’s cancer resistance^18^. Moreover, the authors reported that it was possible the ultrahigh molecular weight form of HA found in the NMR contributed to the elasticity of its characteristic wrinkly skin. With such unusual functions attributed to NMR HA it raises the question of if there is anything in its structure or soft matter properties that could explain its functionality?

The majority of data on the structure/assembly behaviour of HA almost exclusively come from material that is obtained from the fermentation of bacteria (commercially available)^19,20^. Less commonly it is sourced from rooster combs or human fluids and tissues. Whether imaged under aqueous conditions or in air, HA is found to form planar branched networks of fibres^21^. Scott et al. used rotary shadow electron microscopy to show that HA (extracted from rooster combs, *Streptococci* and mesothelioma fluid) forms planar networks in solution, and that the longer the HA molecule, the more branching occurs^22^. More recent studies have used atomic force microscopy (AFM) in air (as opposed to in aqueous solution) to interrogate the morphology of molecular scale nanostructures. Both loose coils of individual molecular chains and extended structures were observed to assemble, with characteristic planar branched network structures common^19,21,23,24^. Using AFM-based force spectroscopy it has been possible to uncoil and unstretch single HA chains and confirm network structures^25^. At the macroscopic scale and in isolation, HA only forms a weak gel and needs to be either chemically crosslinked or combined in a composite to be useful as a biomaterial^26^.

It is well established that HA structural changes over time are fundamental to the ageing of human skin, with its affinity for water thought to be critical for maintaining skin elasticty^27^. To date there has been no attempt to look at the structure or soft matter properties of HA from NMR. Such an understanding of the material properties of NMR HA are a prerequisite for its development as a potential biomaterial for the treatment of cancer. Therefore, the aim of this work was to purify NMR-HA and examine the material and structural properties with particular emphasis on differences and similarities between HA from other species.

## Results and Discussion

The first tissue examined was skin as the characteristic wrinkly, yet stretchy skin of the NMR has been speculatively attributed to the presence of a high molecular weight HA^18^. Using a biotinylated version of the very specific and tightly binding hyaluronan binding protein (HABP)^29^, histological analysis of the plantar surface, forepaw skin demonstrated that HA was present in both mouse and NMR skin. However, HA was observed to a much greater extent in the NMR epidermis compared to the mouse where, as others have shown^30^, HA was largely confined to the dermis (Fig. 1A and 1B). HA extracted from the culture medium of immortalised NMR skin fibroblasts presented a range of assembled conformations, whereby individual chains could clearly be resolved (Figure 1 C and D).

**Fig. 1.**
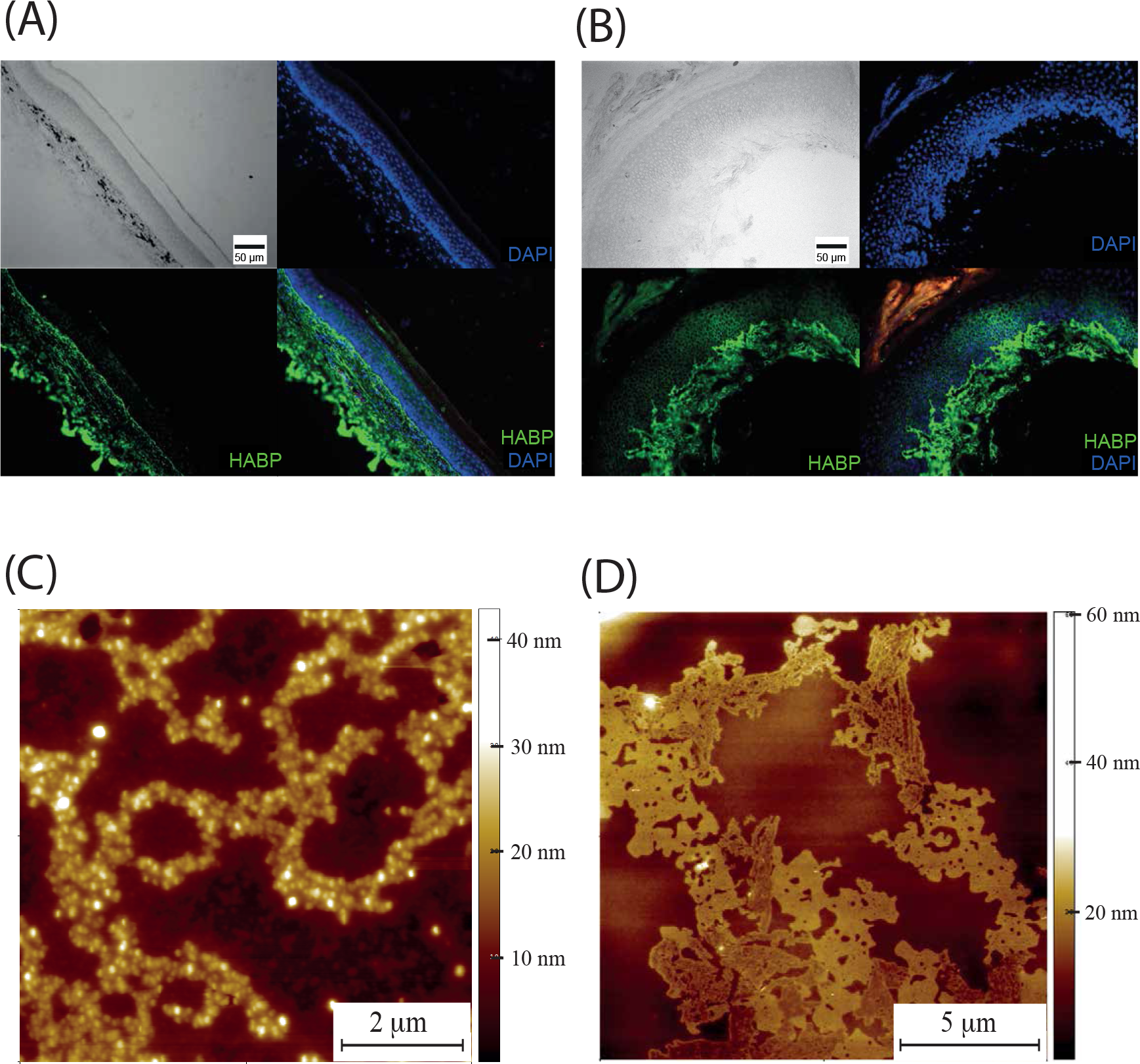
NMR skin HA. (A) Mouse forepaw, plantar skin section stained for HA using HABP and streptavidin-Alexa488 (green) and DAPI nuclear stain (blue). (B) NMR forepaw, plantar skin section stained for HA using HABP and streptavidin-Alexa488 (green) and DAPI nuclear stain (blue). Unlike in the mouse, the NMR HA extends deep into the epidermis and is significantly more abundant. (C) AFM topography image of HA chains obtained from HA extracted from the media of NMR skin fibroblasts. (D) AFM topography image of another type of HA chain assembly from NMR skin fibroblasts.

Dimensions of these chains were consistent with a bundle of a few polymer molecules with an average width of 33.6 ± 7.71 nm (n=100) and height several times greater than the couple of nanometres, the characteristic height of individual molecules. The most abundant structures within a sample were well defined, densely packed supercoils (Fig. 2A and 2B). Such densely folded entities appear to be the basic unit of HA in NMR skin with a characteristic “cauliflower” appearance when visualised with the SEM (Fig. 2C). NMR HA extracted from skin tissue (rather than from the medium of cultured skin fibroblasts) also formed the same supercoiled folded structures, although they were generally larger than those of HA extracted from culture medium (Fig. 2D and 2E). Individual chain networks can also be observed for the NMR skin tissue extracted samples (Fig. 2F), demonstrating a consistency between HA extracted from cell media and HA extracted from tissue. Using a conical AFM tip it was possible to address individual supercoils and determine their soft matter properties via indentation. Young’s modulus values could be obtained by fitting the indentation curve with the Sneddon model, eq 1

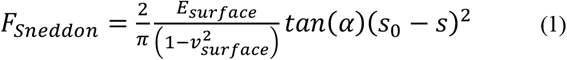

where:

F_Sneddon_ = force
E_surface_ = Young’s modulus of the surface
v_surface_ = Surface Poisson’s ratio
α = Tip half cone opening angle
s_0_ = point of zero indentation
s_0_-s = indentation depth

**Fig. 2.**
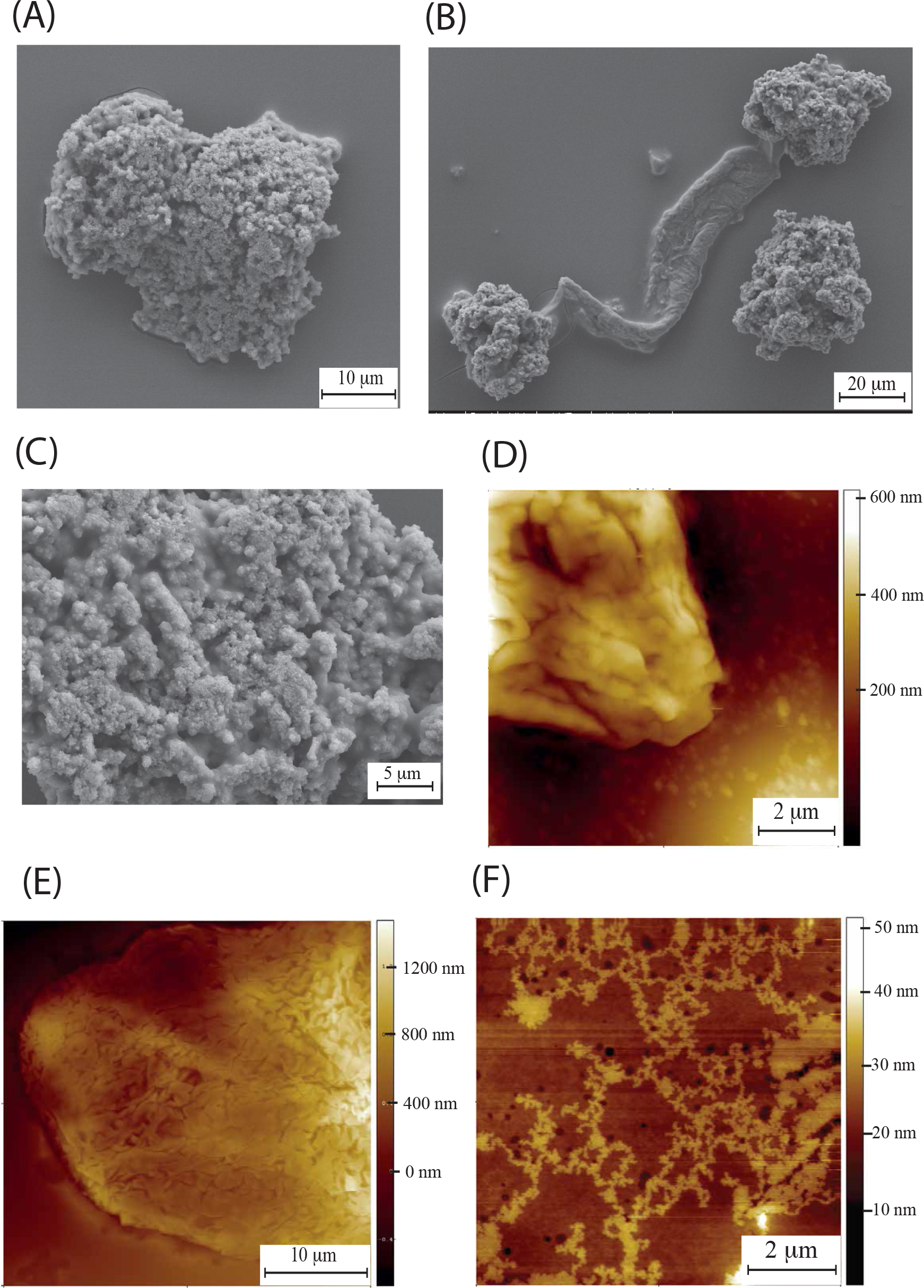
NMR skin HA. (A) Scanning electron microscopy (SEM) of NMR HA extracted from NMR skin fibroblast media.(B) Supercoils linked together with a large fibre. (C) High resolution SEM showing high density of folds (D) AFM of HA extracted from NMR skin fibroblast media (E) AFM of NMR HA extracted from skin tissue. exhibiting again highly folded as in the case of HA extracted from NMR fibroblasts. (F) AFM of individual HA chains extracted from NMR skin tissue.

Fig. 3A shows an AFM topographic image of an individual NMRHA supercoil, with Fig. 3B presenting the individual positions within the supercoil where individual indentation measurements were taken (green squares). A typical force - separation curve for NMR HA extracted from skin is presented in Fig. 3C. Of particular note are the peaks in the red retract curve that represent the unfolding of polymer chains. This is not a uniform sawtooth pattern, but rather an irregularity in terms of widths of the peaks is observed, likely reflective of the different sized folded domains. An unusual feature of this pattern is the peaks in the indentation profile, shown in blue, an interpretation of which would be an unfolding of the structure under compression. From these indentation curves from within skin NMR HA supercoils a value of Young’s Modulus was acquired which was 13.77 ± 2.44 MPa (Fig. 3D).

**Fig. 3.**
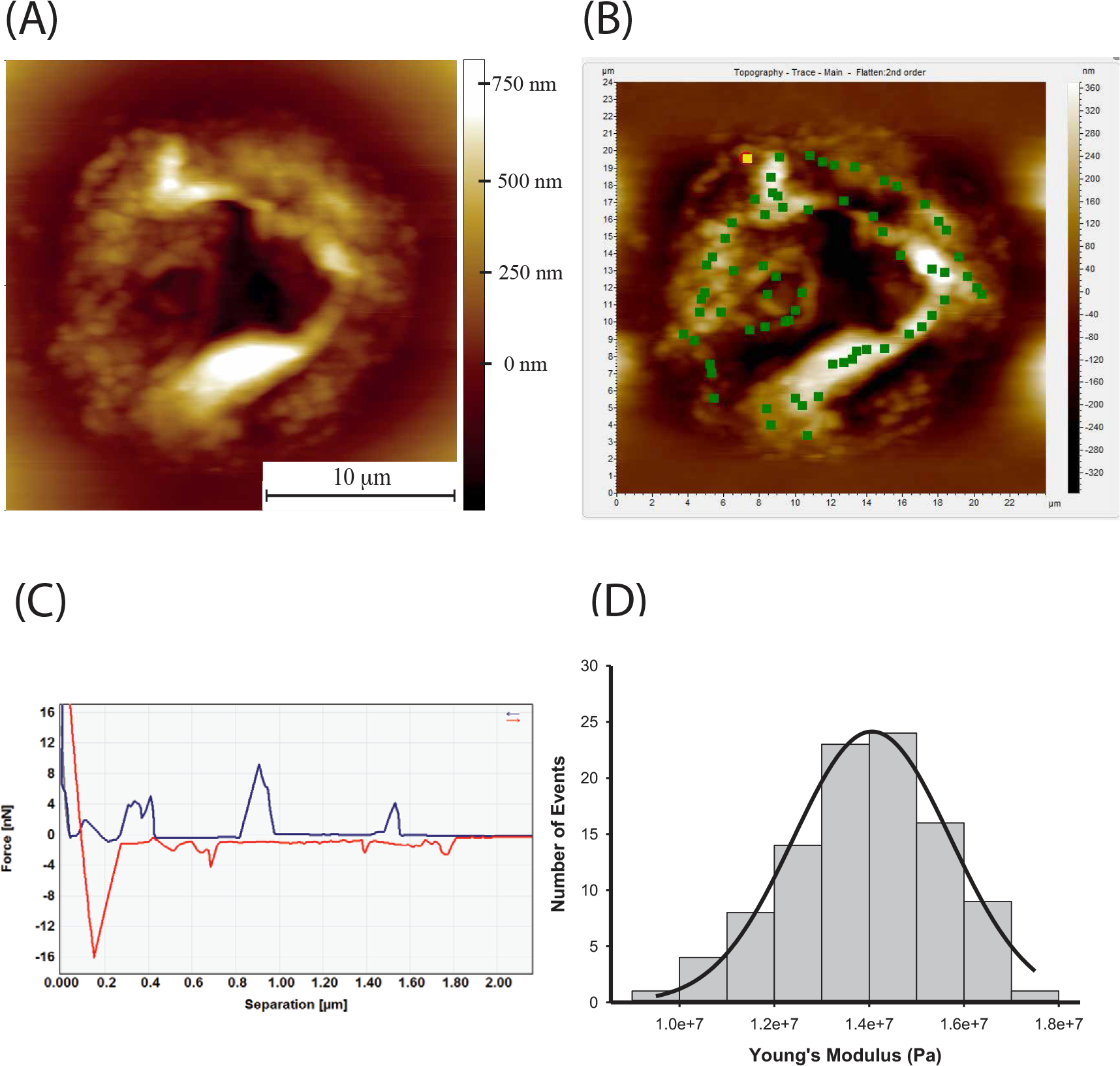
(A) AFM topography image of supercoiled NRM HA structure. These are highly folded structures up to a micron in height and with some folds/twists half a micron in diameter. (B) The same image with a force map superimposed. The green dots indicate where individual indentation measurements are taken. Multiple measurements can be taken at each point and precise control of cantilever is achieved using a closed loop scanner. (C) Force-separation curve (pyramidal tip) for an NMR skin HA supercoil measured in air. The retraction curve (red) of the AFM tip exhibits several peaks over a length of microns suggesting the unfolding of large HA chains. Unusually for a force-separation curve, the approach curve (blue) also exhibits a number of force peaks. We attribute these to the force required to push the supercoiled chains out of the way as the tip penetrates through the structure. (D) Histogram of Young’s modulus values for indentations into NMR skin HA supercoils, taken over 5 coils with 112 curves fitted with the Sneddon model, giving a mean Young’s Modulus value of 13.67 MPa ± 2.44 MPa

As a comparison, HA was extracted and purified from human skin tissue. Unlike the NMR skin HA, there were no supercoiled structures present in human skin HA, but instead large, flat networks were observed, as shown with SEM (Fig. 4A) and AFM (Fig. 4B). The indentation profiles were typical of a soft elastic material (Fig. 4C) with no unfolding peaks in either approach or retract curve. By addressing the AFM tip across the network, the Young’s modulus was determined to be 0.82 GPa ± 0.014 GPa, significantly stiffer than the NMR HA in supercoil form.

**Fig. 4.**
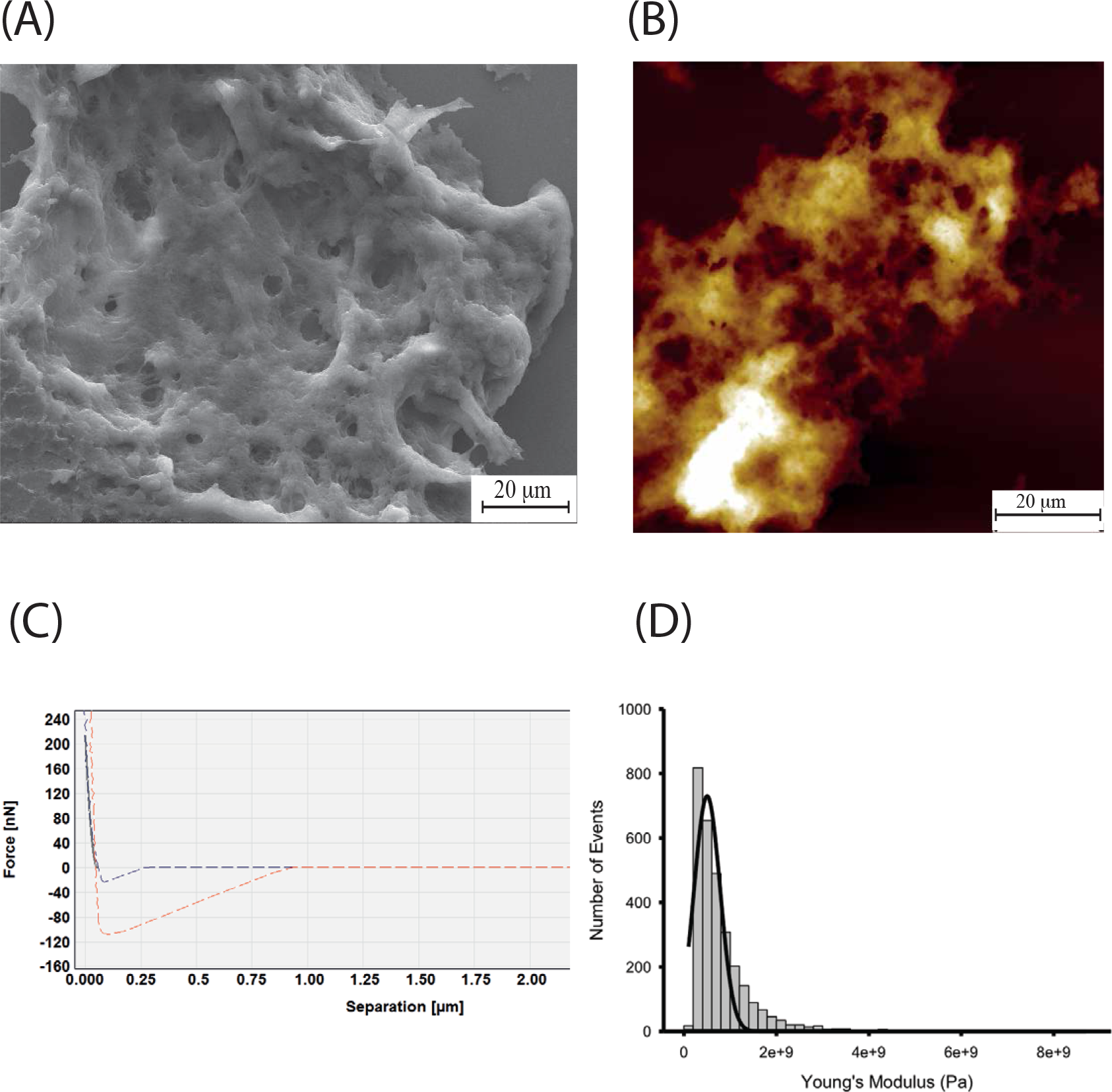
Human skin HA. (A) SEM reveals large planar networks, with well-defined holes up to a couple of microns in diameter. (B) AFM revealing the same form of HA, consistent with the SEM images. (C) Force - separation curve showing indentation into human skin HA network, exhibiting elastic behaviour. (D) Histogram of Young’s modulus values for indentations into human skin HA, taken over large networks. Curves fitted with the Sneddon model, giving a mean Young’s Modulus value of 0.82 GPa ± 0.014 GPa for 3000 curves.

HA was also extracted from NMR brain and lung tissue. HA extracted from whole brain formed large (at least several microns in diameter) supercoils, but this time the folds were less dense and resembled the macroscopic appearance of the gyri and sulci of the human brain (Fig. 5A and 5B). HA extracted from NMR lung also formed supercoils, but this time the characteristic folded structure resembled a morphology which can only be described as resembling a snowman, (Fig. 5C) Higher resolution images revealed these unusual morphologies to be composed of a network of woven chains at high density (Fig. 5D). Both brain and lung HA structures were very tightly folded with few visible gaps. This was in stark contrast to HA extracted from human skin, whereby supercoils were not formed and random networks of chains were the dominant structures. (Commercially available HA (high molecular weight produced by microbial fermentation of *Streptococcus pyogenes*, R&D systems) formed branched networks.

**Fig. 5.**
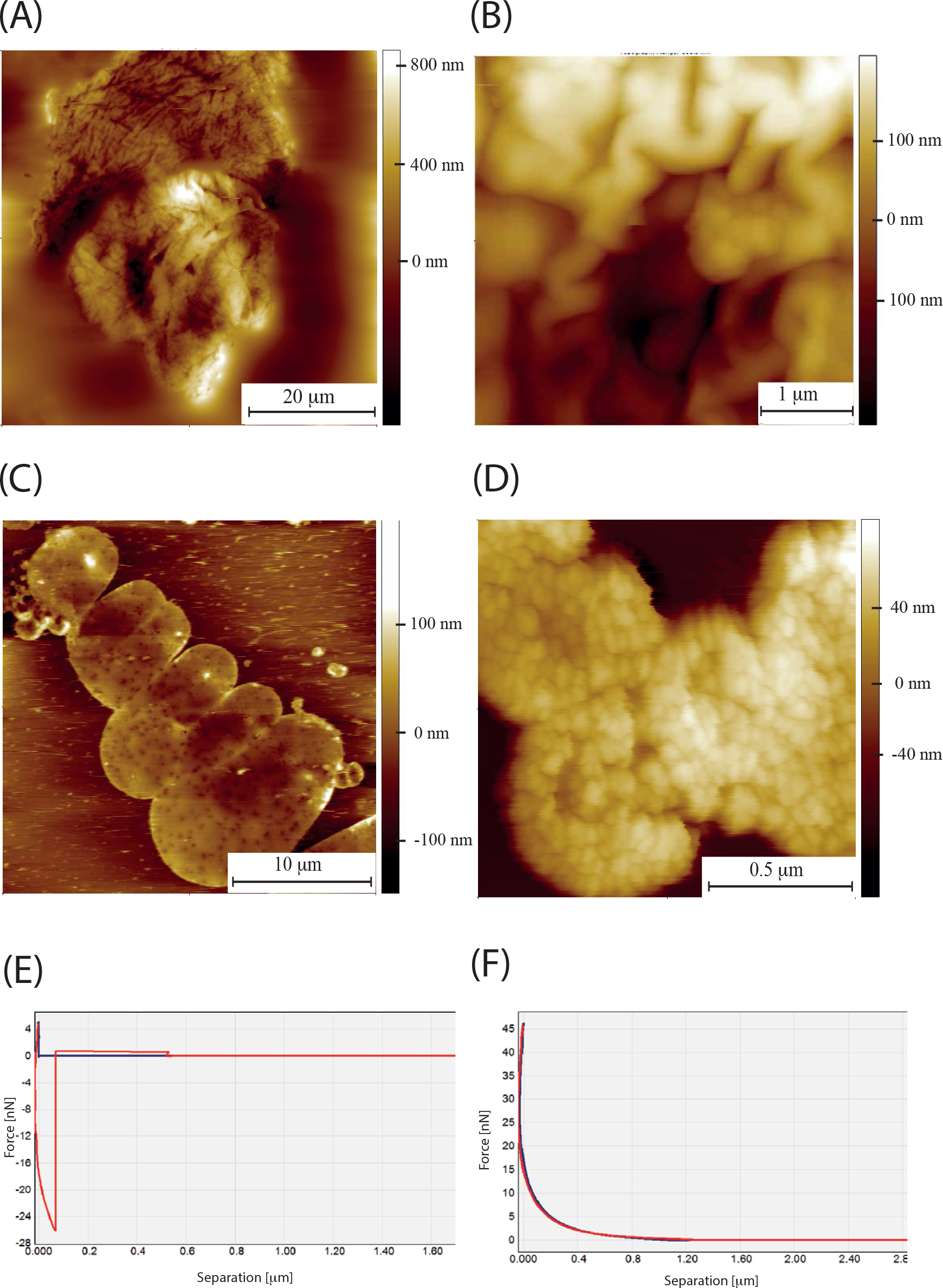
NMR HA extracted from brain and lung. (A) AFM Topographic image of folded NMR HA extracted from whole brain (B) High resolution topographic AFM image exhibiting characteristic “brain-like” folds of HA extracted from NMR whole brain. (C) HA extracted from NMR lung forms characteristic “snowman” structures.(D) Higher resolution scan of HA from NMR lung where the densely folded chains can be resolved, forming an impenetrable network. (E) Indentation curve for NMR lung HA, having been indented multiple times (more than 100) in the same place. The water has been squeezed out and the supercoil now has the characteristics of a purely elastic material. F) The indentation curve for lung NMR HA after having been hydrated for an hour. It has the characteristic form for a hydrogel.

Indentation profiles with peaks in the indentation curve were only seen for the skin supercoils as in Fig. 3C. Upon submerging the lung HA supercoils in water (room temperature) a transition occurs in their mechanical properties. After 10 minutes submerged in water the supercoil retains its elastic characteristics (Fig. 5E), but then after 20 minutes the force-separation curve takes the characteristic form of a viscoelastic material (Fig. 5F). This was the case for HA from all NMR tissues/cells examined. In order to accurately measure the Young’s modulus of supercoils before and after hydration a spherical probe was used. The transition between viscoelastic and elastic, that is from Fig. 5F, to Fig. 5E, can be obtained by repeated indentation into the same position. This repeated loading (sometimes up to 150 indents at a force of up to 50 nN) appears to squeeze out the water. The transition is sudden with the force distance curve changing form between two consecutive indentations. The purely elastic state can also be reached by dehydrating the HA.

For HA extracted from all NMR tissues, that is brain, lung and skin, robust macroscopic gels would spontaneously form by leaving 30 ul of HA solution (HA extract in water) on the mica surface for 15 minutes to dry, leaving a circular gel up to 1 cm in diameter (Fig. 6A). HA extracted from the culture medium of immortalised NMR skin fibroblasts showed the same gelling behaviour, but no such gelling behaviour was observed for HA from mouse or human skin. As the gel dehydrates the constituent microgels are clearly visible (Fig. 6B). The gels have well defined borders (Fig. 6C, D) and the constituent supercoils are recovered upon subsequent dehydration. Indentation curves for the gel particles were fitted with the Hertz model for spheres (Fig. 6E) on a flat surface which assumes that the indent depth is much smaller than the diameter of the probe. Using a spherical probe and fitting the indentation curve to the Hertz model, eq 2, the Young’s modulus of the microgels was determined.

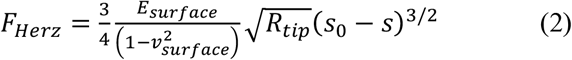

where:

F_Herz_ = force
E_surface_ = Young’s modulus of the surface
v_surface_ = Surface Poisson’s ratio
R_tip_ = ball radius
s_0_ = point of zero indentation
s_0_-s = indentation depth

These were 10.208 ± 1.677 kPa, 10.435 ± 1.312 kPa, 7.862 ± 0.835 kPa, for NMR skin, brain and lung tissue respectively. A typical modulus distribution taken from close to 1000 indentation curves is shown in Fig. 6F. Elastic supercoils (after dehydration) have far superior mechanical properties than the micro gels, for example NMR brain supercoils have a Young’s modulus of 30.305 ± 1.541 MPa (n =1499).

**Fig. 6.**
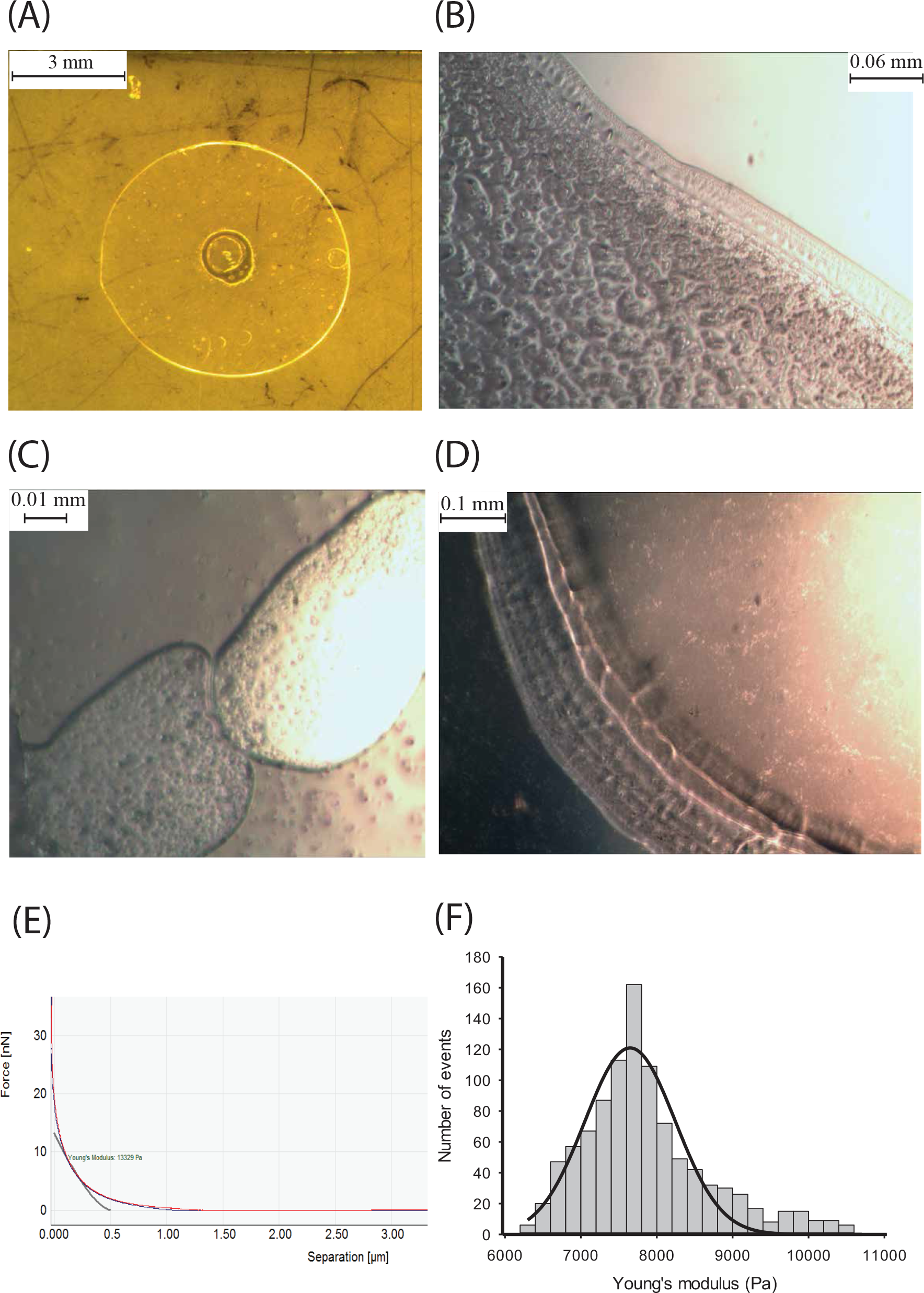
(A) Macroscopic gel formed from NMR HA extracted from skin tissue. (B) Edge of macroscopic gel formed from brain NMR HA. (C) NMR HA gel particles from (derived from brain tissue) (D) Edge of macroscopic gel formed from NMR lung HA. (E) Indentation curve of gel made from NMR skin HA. Spherical probes were used to make these measurements and the Hertz model for a sphere indenting into a surface was fitted to the indentation curve to extract the Young’s modulus. (F) Histogram of Young’s modulus values for indentations into NMR lung HA gel particles using spherical probes. Mean Young’s Modulus value of 7.86 kPa ± 0.834 kPa for 999 curves.

Mouse HA (derived from the medium of skin fibroblasts) does not form supercoils, although less folded spherical particles do exist (Fig. 7A). The most common morphology observed are dense, large scale networks where individual fibres are unable to be resolved (Fig. 7B and 7C) Individual fibres are more abundant for the mouse specimens (Fig. 7D and 7E).

**Fig. 7.**
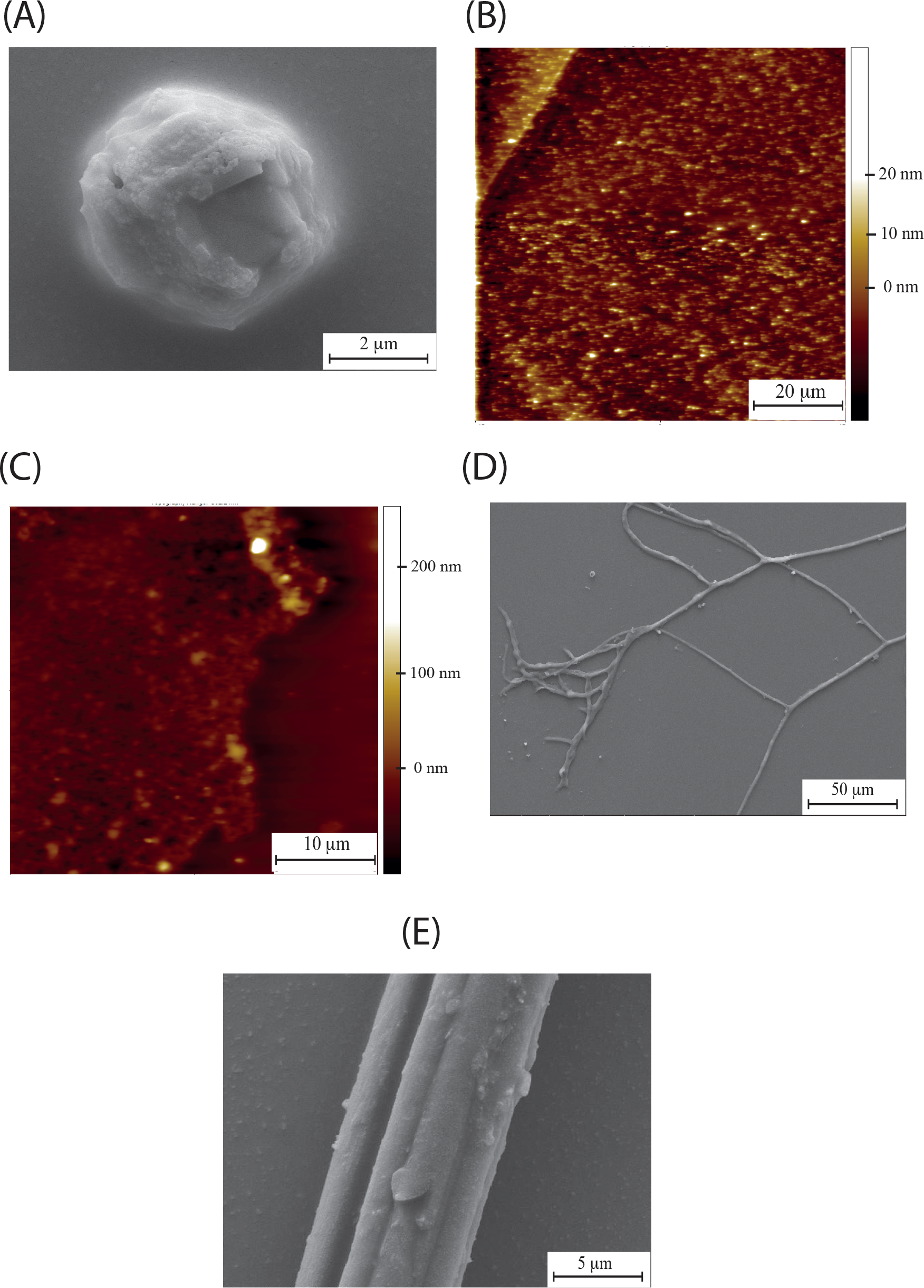
HA extracted from the medium of mouse skin fibroblasts. (A) SEM showing typical spherical particle without a supercoiled structure. SEM showing mouse HA fibres which are more likely to form than for NMR samples. (C) and (D) AFM topographic images of mouse HA random networks. (E) High resolution SEM of mouse HA fibre.

Although supercoils are the dominant morphology for NMR HA (Fig. 8A and 8B), fibres occasionally form. The mechanical properties determined by indentation for the rare fibre that can be found (Fig. 8C) are stiffer than the supercoils, although mechanical heterogeneity along the fibre is clear from the distribution of moduli values. Mouse HA fibres appear to be softer, but absolute values of modulus values using conical tips can be unreliable due to uncertainties in the contact area between tip and sample. What is clear is that NMR fibres are stiffer than the supercoils.

**Fig. 8.**
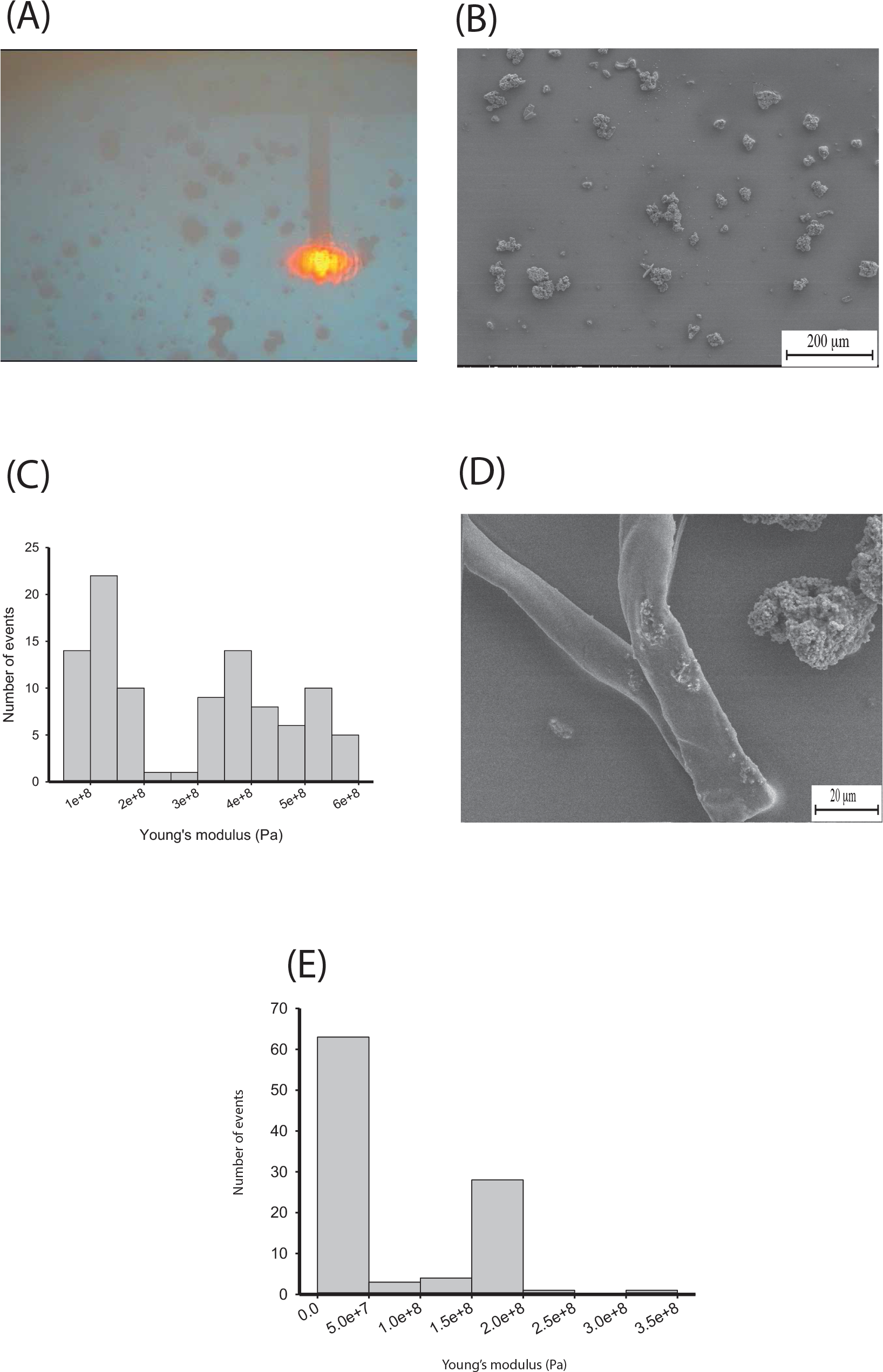
(A) AFM cantilever addressing NMR HA supercoils (B) Large field SEM showing abundance/surface coverage of NMR HA supercoils (B) Histogram of Young’s modulus values taken from a NMR HA fibre with a mean value of 0.28 GPa ± 0.0165 GPa for 100 curves. (D) SEM of NMR HA fibre (E) Histogram of Young’s modulus values taken from a mouse HA fibre with a mean value of 78.025 MPa ± 7.533 MPa for 100 curves.

At higher concentrations adsorbed NMRHA covers the surface and forms a thin film, up to several microns in thickness. Occasionally, the film folds up on itself, presenting the opportunity to observe its internal structure (Fig. 9A) and to determine its thickness of up to 1 micron. At higher resolution a lamellar structure, that is regularly spaced sheets, in this case stacked in pairs, are resolvable (Fig. 9B). The structure within each stack comprises of repeatedly folded chains (Fig. 9C) of similar dimensions to those observed in the supercoiled structures.

**Fig. 9.**
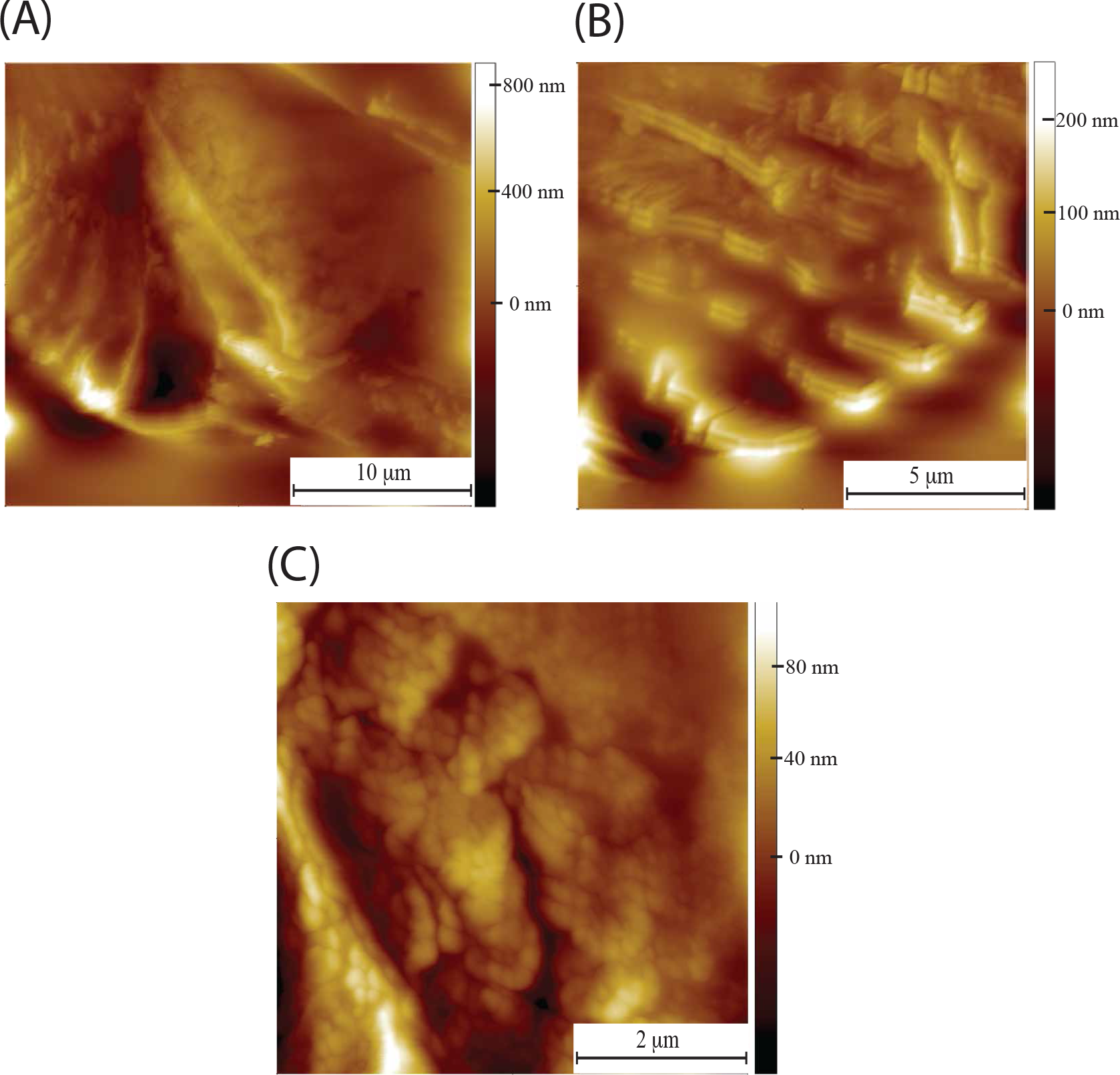
Naked mole rat hyaluronan thin film (A) AFM topographic image showing thin film folded up on itself. The thickness of the film is revealed to be of order of a micron. (B) AFM topographic image presenting the lamellar structure of the thin film. Ledges composed of ordered polymer chains are stacked in pairs each with similar inter planar spacings and morphology. (C) Chains folded along the lamellar of a NMR HA thin film.

## Conclusions

The inability of commercially available HA to spontaneously form robust gels is extensively reported in the literature, with chemical crosslinking or incorporation into a composite being a necessity for practical use as a biomaterial^26,31,32^. By contrast, the nature of NMR HA folding is unusual. It forms a conformation similar to a ball of rubber bands and for the case of HA extracted from skin this leads to a unique indentation profile. Moreover, although the exact concentration could not be determined for such small amounts of biopolymer, the ability of NMR HA to form gels without the need for chemical crosslinking is interesting. A large body of work has been performed on the indentation and unfolding on soft biological matter including proteins, fibres such as collagen, cell membranes, lipid bilayers, and carbohydrates. However, it is rare to find a material that has a sawtooth in an indentation curve (as opposed to a retraction curve).

One example where this does occur is in amyloid fibrils found in natural adhesives. Such an ability to deform elastically in multiple directions would be advantageous for a material that is subjected to shear forces such as those experienced by the skin whilst an animal is squeezing up and round littermates in tunnels. Given the presence of relatively high quantities of HA in NMR skin (Fig. 1B)^18^, it is plausible that these supercoils contribute to its elastic nature. These supercoils with their characteristic cauliflower-like structure are absent from HA extracted from human and mouse skin, as well as being absent from commercially available, bacterially produced HA. In terms of HA structure-function relationships it is well established that the molecular weight of HA can relate to function in the tissue of origin^33^.

For the case of the NMR, it is likely that the ultrahigh molecular weight contributes via supercoiling to the elastic properties of the skin. In addition, the increased surface area of exposed HA will improve water retaining ability, another essential contributor to the anti-ageing properties of NMR skin. This ability to trap water is evidenced by the formation of microgels with the cross linking usually required for robust gels replaced by self-interactions through entangled polymer; importantly, these microgels do not occur for HA extracted from mouse or human skin. Another important feature of the coiled NMR HA is the density of the polymer network. There are few visible gaps due to the folded packing. This is particularly striking when compared to the morphology of human and mouse HA networks. Mimicking the supercoiled structure of NMR HA in either synthetic or engineered natural polymers would be a strategy to explore in recreating the elasticity and anti-ageing effects in tissue engineered skin as well as biomaterials for preventing cell invasion.

## Acknowledgments

This work was supported by a Cancer Research UK Multidisciplinary Project Award (C56829/A22053) to KR, EStJS and DF and a Cancer Research UK Career Establishment Award (C47525/A17348) to WTK. FH and SC were supported by Gates Cambridge Trust scholarships.

## Author Contributions

Ewan St. John Smith, Kenneth Rankin and Daniel Frankel, conceived the study, analysed data and wrote manuscript. Yavuz Kulaberoglu, Bharat Bhushan and Daniel Frankel carried out experiments extracting and characterising HA. Sampurna Chakrabarti carried out immunohistochemistry experiments. Walid Khaled and Fazal Hadi were responsible for cell line construction.

## Materials and Methods

### Animals

All animal experiments were conducted in accordance with the United Kingdom Animal (Scientific Procedures) Act 1986 Amendment Regulations 2012 under a Project License (70/7705) granted to E. St. J. S. by the Home Office; the University of Cambridge Animal Welfare Ethical Review Body also approved procedures. Young adult NMR (both male and female) were used in this study. Animals were maintained in a custom-made caging system with conventional mouse/rat cages connected by different lengths of tunnel. Bedding and nesting material were provided along with a running wheel. The room was warmed to 28 °C, with a heat cable to provide extra warmth running under 2-3 cages, and red lighting (08:00 – 16:00) was used. NMR were killed by exposure to a rising concentration of CO_2_ followed by decapitation. C57Bl6J mice were housed in a temperature controlled (21 °C) room on a 12-hour light/dark cycle, with access to food and water *ad libitum* in groups of up to five. Female mice not older than 12 weeks was used for tissue isolation.

### Immunohistochemistry of NMR and mouse skin samples

Skin samples were collected from the footpad of 6-8 week old female mice (n = 2) and NMR (n = 2), post fixed in 4% paraformaldehyde (PFA) for 1 hr and incubated overnight at 4 °C in 30% sucrose (w/v) for cryo-protection. The skin samples were embedded (epidermal side up) in Shandon M-1 Embedding Matrix (Thermo Fisher Scientific), snap frozen in 2-methylbutane (Honeywell International) on dry ice and stored at −80 °C until further processing. On the day of immunostaining, the embedded skin samples were cut into 12 ◻m sections using a Leica Cryostat (CM3000; Nussloch, Germany) and mounted on Superfrost Plus slides (Thermo Fisher Scientific). For immunohistochemistry, randomly picked slides were washed two times with PBS-Tween, then blocked with an antibody diluent solution, made up of 0.2% (v/v) Triton X-100 and 2% (v/v) bovine serum albumin in PBS, for two hours at room temperature (~22 °C). Biotinylated hyaluronan binding protein (b-HABP) antibody (amsbio, AMS.HKD-BC41, 1:200 in antibody diluent) was added to the slides and incubated overnight at 4 °C. The next day, slides were washed three times using PBS-Tween and incubated for two hours at room temperature with Alexa Fluor 488 Streptavidin (Invitrogen, S11223, 1:1000 in PBS). After the binding of biotin to streptavidin was complete, slides were washed three times in PBS-Tween, and incubated with the nuclear dye, DAPI (Sigma, D9452, 1:1000 in PBS), for 10 min. Slides were further washed once with PBS-Tween, mounted and imaged with an Olympus BX51 microscope (Tokyo, Japan) and QImaging camera (Surrey, Canada). All slides were imaged with the same exposure levels at each wavelengths (488 nm for b-HABP and 350 nm for DAPI), and the same contrast enhancements were made to all slides in ImageJ. Slides that were not incubated with streptavidin conjugate (negative controls) did not show fluorescence.

### Generation of immortalized cell lines

After the animal had been killed, skin was taken from either underarm or underbelly area, cleared of any fat or muscle tissue, generously sprayed with 70% ethanol and finely minced with sterile scalpels. Minced skin was then mixed with 5 ml of NMR Cell Isolation Medium (high glucose DMEM (Gibco #11965092) supplemented with 100 units ml-1 Penicillin, 100μg ml-1 and Streptomycin (Gibco # 15140122) containing 500μl of Cell Dissociation Enzyme Mix (10mg ml^−1^ Collagenase (Roche # 11088793001), 1000Units ml^−1^ Hyaluronidase (Sigma # H3506) in DMEM high glucose (Gibco # 11965092)) and incubated at 37 °C for 3 – 5 hours. Skin was briefly vortexed every 30 minutes to aid cell dissociation and manually inspected for cell dissociation. After complete dissociation, cells were pelleted by centrifuging at 500 g for 5 minutes and resuspended in NMR Cell Culture Medium (DMEM high glucose (Gibco # 11965092) supplemented with 15% fetal bovine serum (Gibco), non-essential amino acids (Gibco # 11140050), 1mM sodium pyruvate (Gibco # 11360039), 100units ml^−1^ Penicillin, 100μg ml^−1^ Streptomycin (Gibco # 15140122) and 100μg ml ^−1^ Primocin (InvivoGen # ant-pm-2)), at 32 °C, 5% CO_2_, 3% O_2_ except for mouse cells which were incubated at 37 °C, 5% CO_2_. To immortalize NMR skin fibroblasts, a lentiviral plasmid carrying SV40LT (SV40LT-hRASg12v was used for mouse cells) was packaged using HEK293FTs as packaging cells. The supernatant of HEK293FT packaging cells was collected, containing SV40 LT-packaged viral particles. When primary NMR skin cells reached 40% confluency, the supernatant containing SV40-packaged viral particles was added on to the primary NMR skin cells. 48 hours after adding the supernatant, cells were treated with 2 μg/ml of puromycin to kill off uninfected cells. Stable immortalized cells were maintained in DMEM supplemented with 15% fetal bovine serum, non-essential amino acids, sodium pyruvate, 100 units ml^− 1^ penicillin, and 100 mg ml^−1^ streptomycin.

### HA extraction from NMR tissues

Following decapitation, tissues were removed, weighed and stored at −80 °C until HA extraction was conducted. Tissues were digested at 50 °C overnight in the digestion buffer (10-mM Tris-CL, 25 mM EDTA, 100 mM NaCI, 0.5% SDS and 0.1 mg/ml of Proteinase K). The next day, samples were centrifuged at 18,000 × g for 10 minutes to remove tissue residual particles and supernatant were transferred to a new tube. Four volumes of ethanol were added into the supernatant, followed by incubation overnight at −20 °C. The following day, samples were cold-centrifuged at 18,000 × g for 10 minutes. Pellets were then washed in four volumes of 75% ethanol by centrifugation. The supernatant was discarded and pellets were incubated at room temperature for 20 minutes to remove any residual ethanol. The pellets were then resuspended in 200 μl of autoclaved distilled water. 500 units of benzonase endonuclease was added and then incubated overnight at 37 °C to remove nucleic acids. The next day, samples were precipitated overnight with one volume of 100% ethanol. The following day, after cold centrifugation at 18,000 × g for 10 minutes, pellets were resuspended in 400 μl of autoclaved distilled water.

### HA extraction from conditioned medium

Immortalized skin fibroblasts were seeded and maintained for 7 days without changing medium. After 7 days, the media from each flask was collected and centrifuged at 4,000 × g for 10 minutes to remove residual cells. After centrifugation, 2 ml of conditioned media were incubated overnight with 500 μg of proteinase K at 50 °C to remove proteins. The following day, 2 volumes of 100% ethanol were added to samples for precipitation and incubated overnight at −20 °C. The next day, samples were cold-centrifuged at 18,000 × g for 10 minutes. After centrifugation, supernatant were discarded and pellets were incubated at room temperature for 20 minutes to allow any residual ethanol to evaporate. After the incubation, the pellet was resuspended in 400 μl of autoclaved distilled water.

### HA extraction from human skin

Following consent to the Newcastle Academic Health Partner’s Bioresource at Newcastle University, skin samples were obtained from patients undergoing orthopaedic surgery. Ethics approval was gained from the REC North East Newcastle 1, Reference Number: REC 12/NE/0395. Human skin was chopped and weighed amount of skin tissues were digested overnight at 50 °C in the digestion buffer (10 mM Tris-Cl, 25 mM EDTA, 100 mM NaCl, 0.5 % SDS and 0.1 mg ml^−1^ proteinase K). After centrifugation, a clear supernatant obtained was mixed with four volumes of prechilled ethanol and incubated at −20 °C for overnight. The precipitate was centrifuged, washed with ethanol and air dried. The pellet was resuspended in 100 mM ammonium acetate solution and incubated with benzonase endonuclease at 37 °C for overnight to remove nucleic acid. The solution was mixed with four volumes of ethanol and incubated at −20 °C for overnight. The pellet obtained after centrifugation was again washed, air dried and resuspended in 100 mM ammonium acetate for further use.

### HA adsorption onto mica

30 ul of purified NMRHA in water was deposited on freshly cleaved muscovite mica and allowed to dry in air for 45 minutes. This is the method reported by Cowman et al for the AFM imaging of HA^24^. They noted that HA adsorbs weakly to mica and cannot be examined under liquid conditions due to HA’s affinity to be in solution. For examining hydrated HA the sample was allowed to dry for 15 minutes so that visible water had evaporated but the sample was still wet.

### Atomic force microscopy

All images were obtained on an Agilent 5500 microscope equipped with close loop scanners. Contact mode imaging was employed for topographic imaging using silicon tips having a nominal force constant of between 0.02-0.77 N/m. Forces were minimized during scanning at a level below 1 nN. Scan rates were between 0.5–1 kHz and all images were recorded at 512 pixel resolution. Measurements were carried out in ultrapure water room at room temperature (~20 °C). Processing and analysis of images was performed using version 6.3.3 of SPIP software (Image Metrology, Lyngby, Denmark).

### Atomic force microscopy based nanoindentation

Two different types of tip indentation geometry were used. The first were conical tips with radius of curvature less than 15 nm. These were used to look at unfolding events within a supercoil. Accurate determination of the spring constant was obtained using the using the equipartition theorem as proposed by Hutter and Bechhoefer^28^. Inaccuracies in terms of knowing the contact area between tip and sample in the Sneddon model for conical tips can be compensated for by using a large spherical probe therefore spherical probes were used to determine Young’s modulus values. Borosilicate spherical probes 5 μm in diameter (NovaScan Technologies) were used to indent supercoils, both wet and dry. Spring constants were determined using the thermal K method.

